# Stability and learning in excitatory synapses by nonlinear inhibitory plasticity

**DOI:** 10.1101/2022.03.28.486052

**Authors:** Christoph Miehl, Julijana Gjorgjieva

## Abstract

Synaptic changes underlie learning and memory formation in the brain. But synaptic plasticity of excitatory synapses on its own is unstable, leading to unlimited growth of synaptic strengths without additional homeostatic mechanisms. To control excitatory synaptic strengths we propose a novel form of synaptic plasticity at inhibitory synapses. We identify two key features of inhibitory plasticity, dominance of inhibition over excitation and a nonlinear dependence on the firing rate of postsynaptic excitatory neurons whereby inhibitory synaptic strengths change in the same direction as excitatory synaptic strengths. We demonstrate that the stable synaptic strengths realized by this novel inhibitory plasticity achieve a fixed excitatory/inhibitory set-point in agreement with experimental results. Applying a disinhibitory signal can gate plasticity and lead to the generation of receptive fields and strong bidirectional connectivity in a recurrent network. Hence, a novel form of nonlinear inhibitory plasticity can simultaneously stabilize excitatory synaptic strengths and enable learning upon disinhibition.

## Introduction

Learning and memory formation in the brain are implemented by synaptic changes undergoing Hebbian plasticity whereby joint pre- and postsynaptic activity increase the strength of synaptic connections (Hebb, 1949; Abbott and Nelson, 2000). However, Hebbian long-term plasticity of excitatory synapses to other excitatory neurons, referred to as excitatory plasticity, is inherently unstable (Miller and MacKay, 1994). Increasing excitatory synaptic strengths leads to an increase in the firing rates of excitatory postsynaptic neurons which in turn further increases synaptic strengths. This positive feedback loop is called ‘Hebbian runaway dynamics’ (Turrigiano and Nelson, 2004). To counteract unstable synaptic growth and control resultant rate dynamics, some form of homeostatic control is needed. Experimental studies have uncovered multiple homeostatic mechanisms. One prominent mechanism is synaptic scaling, where synaptic connections onto a given excitatory neuron potentiate or depress, while preserving relative strengths, to maintain a target level of activity (Turrigiano et al., 1998; Turrigiano, 2008). An alternative mechanism that has gained much recent attention is heterosynaptic plasticity (Lynch et al., 1977; Chistiakova et al., 2015), which occurs both at excitatory and inhibitory synapses that have not been directly affected by the induction of plasticity (Field et al., 2020). A third plausible homeostatic mechanism with significant experimental evidence is intrinsic plasticity which affects the intrinsic excitability of single neurons by adjusting the distribution of different ion channel subtypes (Desai et al., 1999; Debanne et al., 2019).

Various computational studies have benefited from this plethora of experimental evidence for homeostatic control of firing rates and synaptic strengths, and implemented a range of computational models from purely phenomenological ones to detailed biophysical ones. Some relatively straightforward ways to stabilize firing rates and control synaptic strengths in models include imposing upper bounds on synaptic strengths, applying normalization schemes which adjust synaptic strengths by preserving the total sum of incoming weights into a neuron (Oja, 1982; Miller and MacKay, 1994) and assuming that the plasticity mechanism modifying synaptic strengths is itself plastic – called ‘metaplasticity’ (Bienenstock et al., 1982; Yger and Gilson, 2015). These can often be linked to the above experimentally described homeostatic mechanisms. Computational studies have also begun to uncover the various, often complementary, functional roles of different homeostatic mechanisms, e.g. on synaptic scaling versus intrinsic plasticity (Wu et al., 2020) or heterosynaptic plasticity (Field et al., 2020). However, how exactly synaptic plasticity and homeostatic mechanisms interact to control synaptic strengths, and yet enable learning, is still unresolved (Fox and Stryker, 2017; Turrigiano, 2017; Yee et al., 2017). Part of the challenge is that the experimentally measured timescales of homeostatic mechanisms are too slow to stabilize the Hebbian runaway dynamics in computational models, sometimes referred to as the ‘temporal paradox’ of homeostasis (Zenke et al., 2013, 2017; Zenke and Gerstner, 2017). A related problem to the integration of plasticity and homeostasis is the trade-off between stability and flexibility. While stimulus representations need to be stable, for instance to allow long-term memory storage, the system also needs to be flexible to allow re-learning of the same, or learning of new representations (Fusi, 2017).

Here, we investigate an under-explored mechanism to control and stabilize excitatory synaptic strengths and their dynamics, which is long-term plasticity of inhibitory-to-excitatory (I-to-E) synapses, referred to as inhibitory plasticity. Experimental paradigms have characterized diverse forms of inhibitory plasticity, usually via high-frequency stimulation (Caillard et al., 1999; Shew et al., 2000; Mellor, 2018) and via pairing of presynaptic and postsynaptic spikes (D’amour and Froemke, 2015; Hennequin et al., 2017). Inhibition has been shown to control the plasticity mechanisms regulating connection strengths between excitatory neurons depending on their firing rates (Steele and Mauk, 1999) as well as precise spike timing (Paille et al., 2013; Hiratani and Fukai, 2017; Herstel and Wierenga, 2021). Inhibitory plasticity can even dictate the direction of excitatory plasticity, shifting between depression or potentiation (Wang and Maffei, 2014). Given this potential of inhibitory plasticity to affect so many different aspects of synaptic strength and firing rate dynamics in a network, it remains unclear what properties are important for achieving stability, while still enabling neural circuits to learn.

Using computational modeling, we characterize a novel mechanism of inhibitory plasticity with two key features. First, we propose that inhibitory plasticity should depend nonlinearly on the firing rate of an excitatory postsynaptic neuron to robustly control and stabilize the strengths of excitatory synaptic connections made by that neuron. This means that for low postsynaptic rates, I-to-E synapses should depress, for high postsynaptic rates I-to-E synapses should potentiate and without any postsynaptic activity undergo no plasticity. This nonlinear dependence of inhibitory plasticity on the postsynaptic firing rate is sufficient for stability, without the need for additional homeostatic mechanisms. Second, we require a dominance of inhibition, which can be reflected in the larger number of synaptic connections, faster synaptic dynamics or overall higher firing rates of inhibitory synapses and neurons relative to excitatory ones. Dominance of inhibition has already been demonstrated in circuits in the visual cortex which operate as inhibition-stabilized networks (ISNs) (Tsodyks et al., 1997; Sanzeni et al., 2020; Ahmadian and Miller, 2021). A direct consequence from our proposed novel mechanism of nonlinear inhibitory plasticity is the formation of a fixed ratio of excitatory to inhibitory synaptic strengths, in agreement with experimental data (D’amour and Froemke, 2015). Besides stability, our proposed inhibitory plasticity can also support flexible learning of receptive fields and recurrent network structures by gating excitatory plasticity via disinhibition (Froemke et al., 2007; Letzkus et al., 2011). Therefore, our results provide a plausible solution to the stability-flexibility problem by identifying key aspects of inhibitory plasticity, which provide experimentally testable predictions.

## Results

### A linear inhibitory plasticity rule fails to robustly stabilize weight dynamics

To investigate the plausibility of inhibitory plasticity as a control mechanism of excitatory synaptic strengths, we initially considered a model based on a feedforward inhibitory motif prominent in many brain circuits (Fig. 1A). Here, a population of presynaptic excitatory neurons projects to a population of inhibitory neurons and both populations project to a postsynaptic excitatory neuron. Such a motif could resemble, for instance, the excitatory input from the thalamus to excitatory and inhibitory neurons in a primary sensory cortical area (Tremblay et al., 2016). We described the activity of neurons by their firing rates, and investigated average population firing rates and synaptic strength changes as a function of synaptic plasticity (Methods). Experimental studies have shown that the sign and magnitude of excitatory plasticity depends nonlinearly on the firing rates (Kirkwood et al., 1996; Philpot et al., 2003; Cooper and Bear, 2012). Inspired by these findings, we implemented plasticity of E-to-E synaptic connections *w*^*EE*^ (or weights) as a nonlinear function of the postsynaptic rate *v*^*E*^ (Fig. 1B):

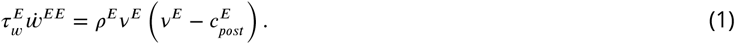

**Figure 1.**
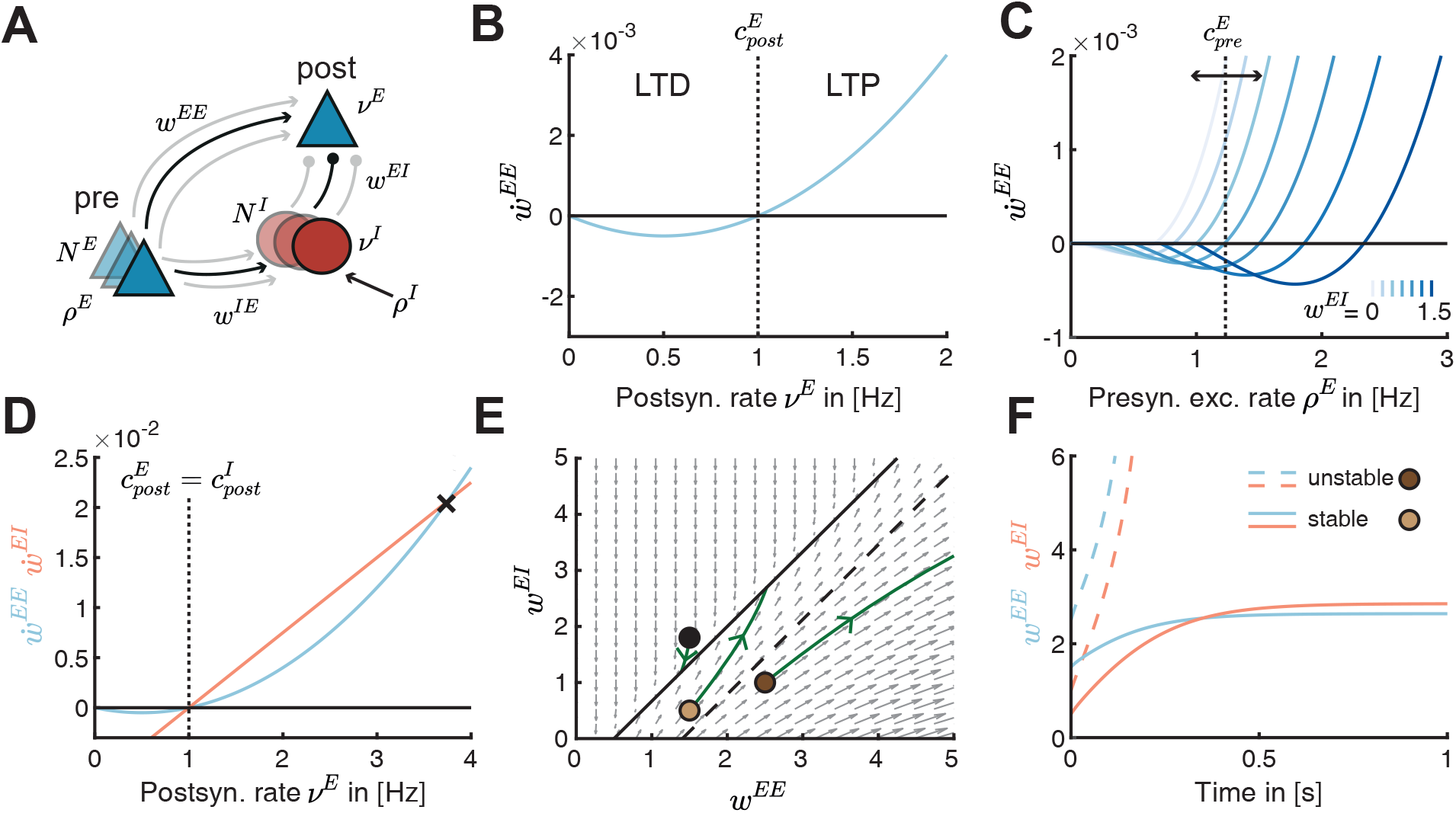
Linear inhibitory plasticity fails to stabilize weights for high excitatory firing rates. **A**. Schematic of a feedforward inhibitory motif. A single postsynaptic excitatory neuron with rate *v*^*E*^ receives input from an excitatory presynaptic population with number of synapses *N*^*E*^, firing rate *ρ*^*E*^ and weight *w*^*EE*^ and an inhibitory neuron population with number of synapses *N*^*I*^, firing rate *v*^*I*^ and weight *w*^*EI*^. The inhibitory population receives external excitatory input with rate *ρ*^*I*^ and input from the presynaptic excitatory population via *w*^*IE*^. **B**. Plasticity curve of E-to-E weights (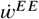, blue) as a function of the postsynaptic rate *v*^*E*^. The postsynaptic LTD/LTP threshold 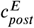 is set to 1 Hz. **C**. E-to-E weight change 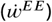 as a function of the presynaptic excitatory rate *ρ*^*E*^ for different I-to-E weights *w*^*EI*^ ranging from 0 to 1.5. The presynaptic LTD/LTP threshold 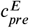 is shown for *w*^*EI*^ = 0.75 (vertical dashed line). **D**. Plasticity curves of E-to-E (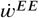, blue) and I-to-E (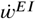, red) weights as a function of the postsynaptic rate *v*^*E*^. The excitatory and inhibitory LTD/LTP thresholds are identical 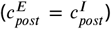. Black cross marks crossover of the plasticity curves at which weight dynamics become unstable. **E**. Phase portrait of the dynamics of E-to-E (*w*^*EE*^) and I-to-E (*w*^*EI*^) weights in the phase plane. Grey arrows indicate the sign of weight evolution over time, points represent three different weight initializations, 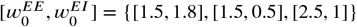, and green lines represent the weight evolution for each case. The two colored points represent initial weights in F. Black line indicates the line attractor and the dashed line separates stable from unstable initial conditions (Methods, Eq. 19). **F**. E-to-E (*w*^*EE*^, blue) and I-to-E (*w*^*EI*^, red) weights as a function of time for stable (solid lines, 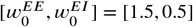) and unstable (dashed lines, 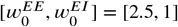) initial conditions.

Here, *ρ*^*E*^ denotes the excitatory presynaptic rate and 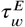 is the timescale of excitatory plasticity. We refer to the postsynaptic rate at which the plasticity changes sign as the ‘postsynaptic LTD/LTP threshold’, denoted by 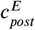. If the firing rate *v*^*E*^ is smaller than the threshold 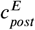, then the change in synaptic strength is negative leading to long-term depression (LTD), while if *v*^*E*^ is larger than 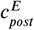, then the change in synaptic strength is positive leading to long-term potentiation (LTP) (Fig. 1B). This means that increasing the excitatory postsynaptic firing rate will lead to potentiation of excitatory weights, and in a positive feedback loop will further increase the neuron’s firing rate – known as the classical problem of ‘Hebbian runaway dynamics’.

Hence, we wanted to determine a plausible mechanism to counteract excitatory runaway dynamics. We postulated that regulating the inhibitory input into the postsynaptic neuron provides an efficient way to stabilize excitatory weights and firing rates. In our framework, inhibitory neurons can affect excitatory plasticity in three equivalent ways. (1) The number of inhibitory synapses *N*^*I*^ onto the postsynaptic neuron can change, for example, through the growth or removal of synapses via structural plasticity. (2) The strength of I-to-E synapses *w*^*EI*^ can change via inhibitory plasticity. (3) Finally, the rate of inhibitory neurons *v*^*I*^ can also change through the external excitatory input to the inhibitory neurons *ρ*^*I*^ or the excitatory-to-inhibitory weight *w*^*IE*^. Various experimental studies have revealed that the plasticity of I-to-E synapses can be induced via the stimulation of the relevant input pathways (Caillard et al., 1999; Shew et al., 2000; Wang and Maffei, 2014). Given this experimental evidence for the plasticity of I-to-E synapses, we examined the influence of changing the strength of I-to-E synapses, *w*^*EI*^, on the strength and magnitude of E-to-E synapses, *w*^*EE*^ (Fig. 1C). We found that stronger *w*^*EI*^ weights rates require higher presynaptic excitatory rates to induce LTP, while weaker *w*^*EI*^ weights require lower presynaptic excitatory rates to induce LTP. This effectively leads to a shift of the threshold between LTD and LTP as a function of the presynaptic excitatory firing rate as *w*^*EI*^ changes. We refer to the presynaptic excitatory firing rate at which the plasticity changes sign between potentiation and depression as the ‘presynaptic LTD/LTP threshold’, denoted by 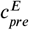 (Fig. 1C). In contrast to the fixed postsynaptic LTD/LTP threshold, 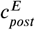 (Fig. 1B), this presynaptic LTD/LTP threshold depends, among others, on the strength of I-to-E synapses (Fig. 1C; Methods, Eq. 12).

Rather than hand-tuning the strength of I-to-E synapses, here we propose that a particular inhibitory plasticity rule can dynamically adjust their strength as a function of presynaptic inhibitory and postsynaptic excitatory activity. However, the exact form of this plasticity has not yet been mapped experimentally. Therefore, we first investigated an inhibitory plasticity rule widely-used in computational models which depends linearly on the post-synaptic rate *v*^*E*^ (Vogels et al., 2011; Clopath et al., 2016) (Fig. 1D, 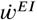):

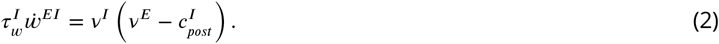

Here, 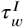 denotes the timescale of inhibitory plasticity. As for excitatory plasticity, we refer to the postsynaptic rate at which inhibitory plasticity changes from LTD to LTP as the ‘inhibitory postsynaptic LTD/LTP threshold’, denoted by 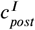. This threshold determined the ‘target rate’ of the postsynaptic neuron (Vogels et al., 2011). If the excitatory postsynaptic neuron fires at higher rates than 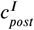, inhibitory LTP leads to a decrease of its firing rate, while if the neuron fires at lower rates than 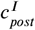, inhibitory LTD increases its rate. To prevent an unstable scenario where excitatory (Eq. 1) and inhibitory plasticity (Eq. 2) push the postsynaptic excitatory neuron towards two different firing rates, here we assume that the excitatory and inhibitory thresholds are matched (Fig. 1D, 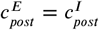).

To investigate the effect of this ‘linear inhibitory plasticity’ mechanism on the temporal evolution of excitatory and inhibitory synaptic weights, *w*^*EE*^ and *w*^*EI*^, we plotted the flow field in the phase plane *w*^*EI*^ vs. *w*^*EE*^ (Fig. 1E). We found that the interaction of excitatory and inhibitory plasticity generates a line of stable fixed points (i.e. a line attractor) where both synaptic weights do not change any more (Fig. 1E, black solid line; see Methods). The initial weights determine whether the weights ultimately converge to the line attractor and stabilize. When the initial E-to-E weights *w*^*EE*^ are much larger than the initial I-to-E weights *w*^*EI*^ (Fig. 1E, below the dashed line), the weights become unstable (Fig. 1E, F). Equivalently, the weights become unstable when the postsynaptic rate *v*^*E*^ is beyond the crossover point of the excitatory and inhibitory plasticity curves as a function of the postsynaptic excitatory rate (Fig. 1D, black cross). For firing rates beyond this crossover point, the E-to-E weights increase faster than the I-to-E weights, leading to runaway dynamics.

In summary, our results suggest that a well-known form of inhibitory plasticity with a linear dependence on the postsynaptic excitatory firing rate can control excitatory weight changes only for a range of initial conditions. There exists a whole range of initial conditions (specifically where the E-to-E are larger than the I-to-E weights) where the postsynaptic excitatory firing rate is sufficiently large and where the weight dynamics explode. This scenario could be problematic if during normal development in the animal, the E-to-E and I-to-E weights are set up in this range, and implies the need for careful tuning to prevent unlimited weight growth.

### A novel nonlinear inhibitory plasticity rule as a robust mechanism to stablize excitatory weights

To ensure weight stability without fine tuning of the initial E-to-E and I-to-E weights, we proposed a novel inhibitory plasticity rule. The rule depends nonlinearly on the postsynaptic rate *v*^*E*^, similarly to excitatory plasticity (Eq. 1, Fig. 2A):

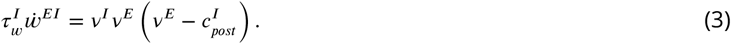

**Figure 2.**
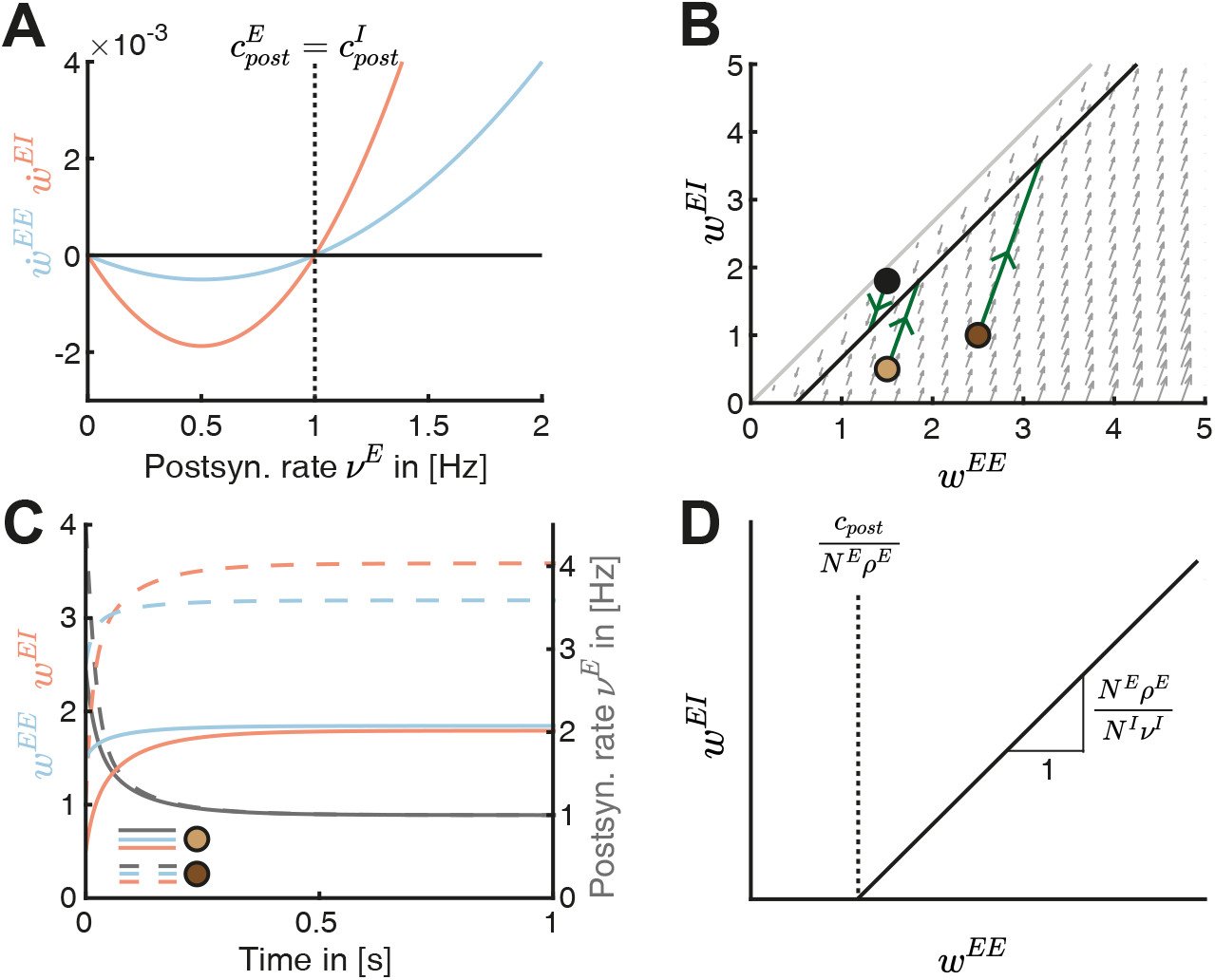
A novel nonlinear inhibitory plasticity rule can counteract runaway dynamics of excitatory-to-excitatory weights. **A**. Plasticity curves of E-to-E (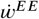, blue) and I-to-E (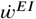, red) weights as a function of the postsynaptic rate *v*^*E*^. The excitatory and inhibitory LTD/LTP thresholds are identical 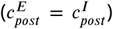. **B**. Phase portrait of the dynamics of E-to-E (*w*^*EE*^) and I-to-E (*w*^*EI*^) weights in the phase plane. Grey arrows indicate the direction of weight evolution over time, points represent three different initial conditions of the weights, 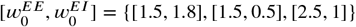, and green lines represent the weight evolution for each initial condition. The two colored points represent initial weights in C. Black line indicates the line attractor and the grey line separates the space at which the postsynaptic firing rate is zero (no dynamics) or larger than zero (Methods, Eq. 17). **C**. E-to-E (*w*^*EE*^, blue) and I-to-E (*w*^*EI*^, red) weight dynamics and postsynaptic rate dynamics (*v*^*E*^, grey) as a function of time for two initial conditions in B, 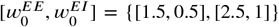. **D**. The slope and intersection of the line attractor with the abscissa (black line) depend on the number and firing rates of excitatory and inhibitory neurons and the LTD/LTP threshold.

As before, to prevent a scenario where the two, excitatory and inhibitory, plasticity rules push the postsynaptic excitatory neuron towards two different firing rates, we assume here that the excitatory and inhibitory thresholds are matched 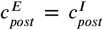. However, as we show later, this assumption can be relaxed. Differently from the linear inhibitory plasticity rule (Eq. 2), the nonlinear inhibitory plasticity rule ensures that I-to-E synapses do not change in the case where the postsynaptic firing rate is zero (Fig. 2B, beyond grey line), as suggested experimentally (Wang and Maffei, 2014). Additionally, the nonlinear rule eliminates the region of initial weight configurations in the phase space where the weights grow out of bound; instead the weights converge to the line attractor (Fig. 2B). Indeed, the E-to-E weights, I-to-E weights and the postsynaptic rate reach a stable configuration over time (Fig. 2C). We calculated the condition leading to stable weight dynamics (Methods, Eq. 13-16) as a function of the excitatory and inhibitory input rates (*v*^*I*^, *ρ*^*E*^), the number of synapses (*N*^*E*^, *N*^*I*^) and the timescale of the plasticity mechanisms 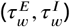:

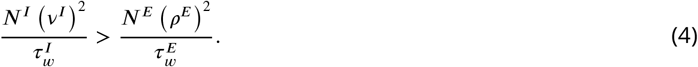

This condition ensures stable weight dynamics whenever inhibition is more ‘dominant’ than excitation, either by having more inhibitory synapses (*N*^*I*^), higher inhibitory rate (*v*^*I*^), a faster timescale of inhibitory plasticity 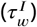 or a combination thereof. From now on, we assume a faster timescale of inhibitory relative to excitatory plasticity (see Table 1). An alternative way to achieve stability involves a feedback connection from the postsynaptic neuron to the inhibitory population (Suppl. Fig. S1A). In this case, sufficiently strong E-to-I feedforward and feedback weights guarantee stability in the presence of this feedback inhibitory motif (Suppl. Fig. S1B-D).

**Table 1.**
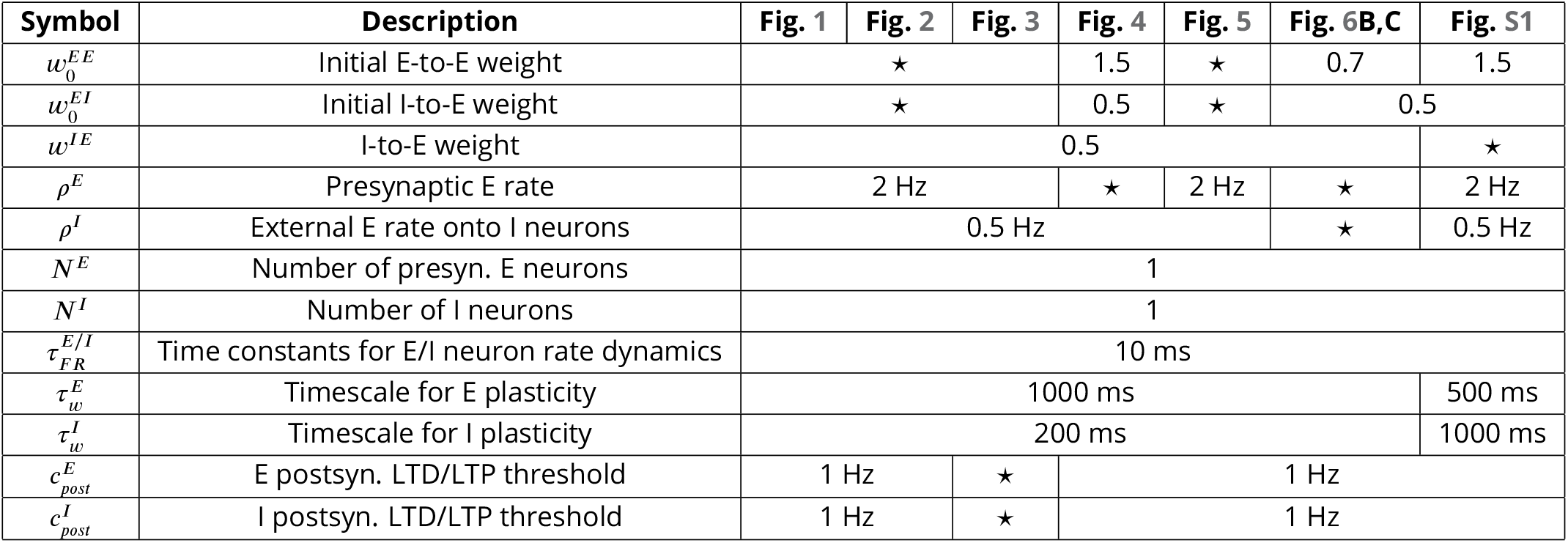
Parameter values for figures, ⋆ denotes that values are provided in the figure captions.

We found that the line attractor depends on several model parameters (see Methods, Eq. 13) (Fig. 2D)

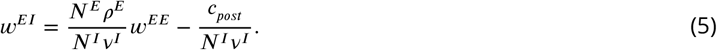

Under the assumption that the LTD/LTP thresholds of excitatory and inhibitory plasticity are the same, 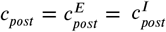, we found that the slope of the line attractor can be written as *N*^*E*^ *ρ*^*E*^ / (*N*^*I*^ *v*^*I*^), while the intersection of the line attractor with the abscissa can be written as *c*_*post*_/ (*N*^*E*^ *ρ*^*E*^). Therefore, by changing any of the network parameters we can predict the stable configuration to which the weights will converge.

Taken together, we have proposed a novel form of nonlinear inhibitory plasticity which can counteract excitatory runaway weight dynamics without the need for fine tuning. The proposed rule eliminates the need for additional homeostatic mechanisms and upper bounds on the weights to stabilize weight dynamics. Our modeling approach allows us to dissect the exact dependencies of the stability condition on number of synapses, firing rates and plasticity timescales of excitatory and inhibitory neurons.

### Dynamic matching of the excitatory and inhibitory postsynaptic thresholds between LTD and LTP

What happens if the postsynaptic thresholds between LTD and LTP for excitatory and inhibitory synapses are not identical, as might be the case in most biological circuits (Fig. 3A)? We found that this leads to the disappearance of the line attractor (see Methods Eq. 13). When the excitatory postsynaptic threshold is lower than the inhibitory postsynaptic threshold 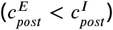, both E-to-E and I-to-E weighs grow unbounded (Fig. 3B). E-to-E weights cannot stabilize as they continue to potentiate 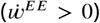 even though the postsynaptic neuron is controlled by the fast inhibitory plasticity and approaches the target rate 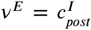 (Fig. 3C). Therefore, stability of firing rates does not imply stability of synaptic weights, especially in the case when the postsynaptic thresholds between LTD and LTP are non-equal. In the case of 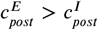, E-to-E and I-to-E weights steadily decrease.

**Figure 3.**
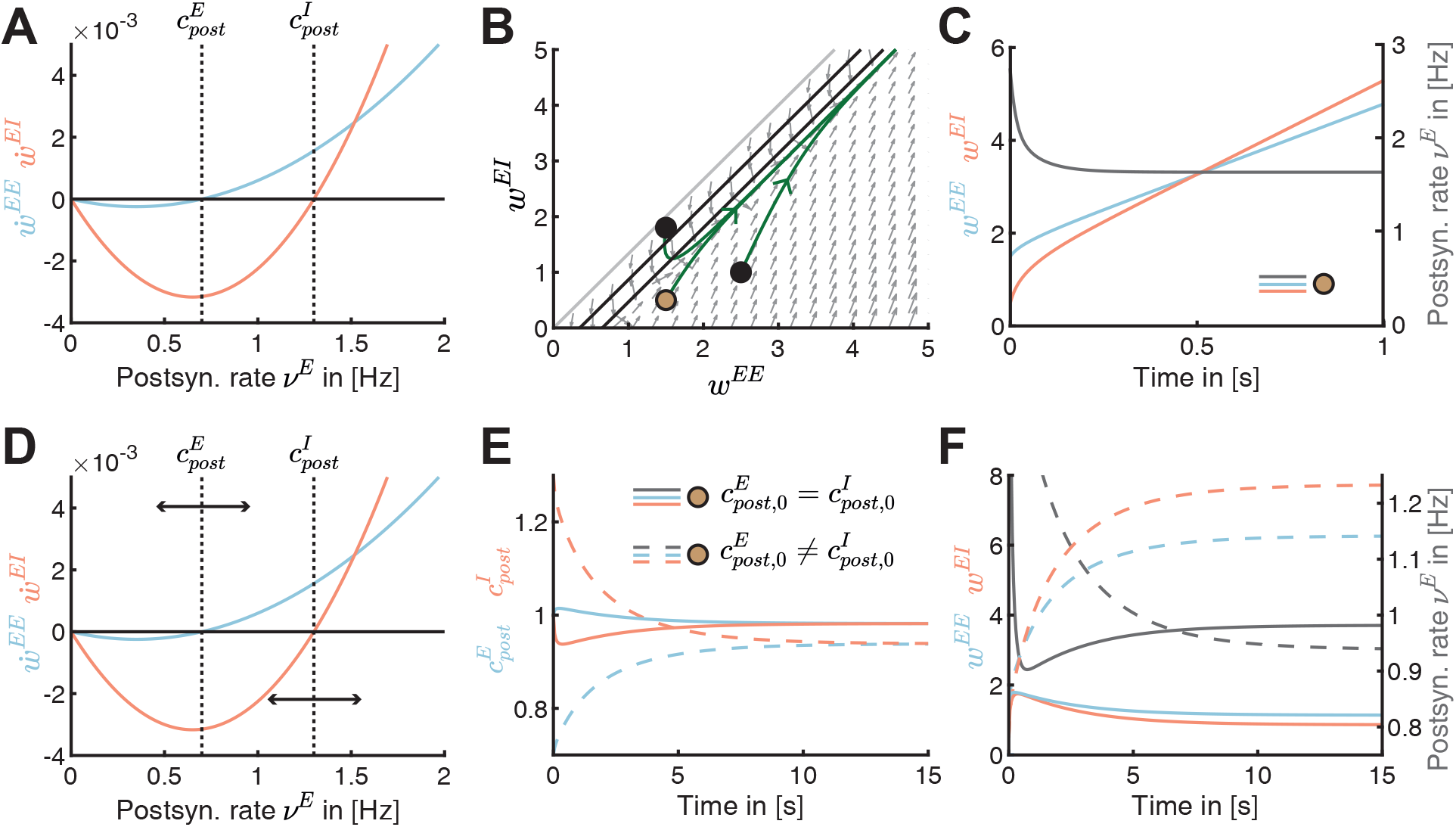
Dynamic matching of the excitatory and inhibitory postsynaptic LTD/LTP thresholds. **A**. Plasticity curves of E-to-E (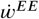, blue) and I-to-E (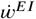, red) weights as a function of the postsynaptic rate *v*^*E*^ with static, non-identical LTD/LTP thresholds 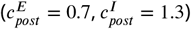. **B**. Phase portrait of the dynamics of E-to-E (*w*^*EE*^) and I-to-E (*w*^*EI*^) weights in the phase plane for the scenario with static thresholds in A. Grey arrows indicate the direction of weight evolution over time, points represent three different initial conditions of the weights, 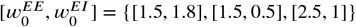, and green lines represent the weight evolution for each initial condition. The colored point represents initial weight in C and E-F. Black lines indicate the nullclines and the grey line separates the space at which the postsynaptic firing rate is zero (no dynamics) or larger than zero (Methods, Eq. 17). **C**. Excitatory (*w*^*EE*^, blue) and inhibitory (*w*^*EI*^, red) weight dynamics and postsynaptic rate dynamics (*v*^*E*^, grey) for one initial condition in B, 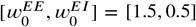. The thresholds are static as in A. **D**. Postsynaptic LTD/LTP thresholds 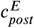 and 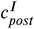 shift dynamically depending on recent postsynaptic rate *v*^*E*^. For lower postsynaptic rate than the excitatory postsynaptic LTD/LTP threshold 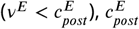 decreases, and for 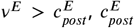 increases. For higher postsynaptic rate than the inhibitory postsynaptic LTD/LTP threshold 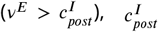 decreases, and for 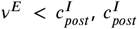 increases (see Methods). **E**. Evolution of excitatory (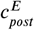, blue) or inhibitory (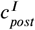, red) postsynaptic LTD/LTP thresholds for two different initial conditions 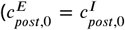, full lines and 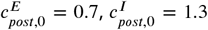, dashed lines). Same initial weight condition as in C, 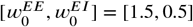, but for dynamic thresholds shown in D. **F**. Excitatory (*w*^*EE*^, blue) and inhibitory (*w*^*EI*^, red) weight dynamics and postsynaptic rate dynamics (*v*^*E*^, grey) for two different initial conditions (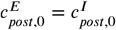, full lines and 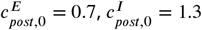, dashed lines). Same initial weight condition as in 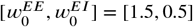, but for dynamic thresholds shown in D. See E for the legend.

Motivated by experimental findings and theoretical considerations (Keck et al., 2017), we proposed that these thresholds can be dynamically regulated in opposite directions (Fig. 3D; see Methods). When the postsynaptic rate is lower than the excitatory postsynaptic LTD/LTP threshold 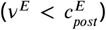, the excitatory postsynaptic LTD/LTP threshold should decrease, while when the postsynaptic rate is higher than the threshold 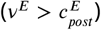, the excitatory threshold should increase. Similarly, when the postsynaptic rate is higher than the inhibitory postsynaptic LTD/LTP threshold 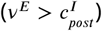, the inhibitory postsynaptic LTD/LTP threshold should decrease, while when the postsynaptic rate is lower than the threshold 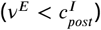, the inhibitory threshold should increase. Eventually, these dynamic lead to matching excitatory and inhibitory LTD/LTP thresholds (Fig. 3E). Therefore, the rate and weight dynamics can both be simultaneously stabilized (Fig. 3F). Implementing this dynamic threshold adjustment process generates different postsynaptic LTD/LTP threshold configurations (Fig. 3E) and postsynaptic rates (Fig. 3F, grey lines). The generation of such heterogeneous postsynaptic rates is consistent with experimental observations in multiple brain regions (Buzsáki and Mizuseki, 2014).

### The nonlinear inhibitory plasticity rule can regulate the network response to perturbations

Since even if they are unequal, excitatory and inhibitory LTD/LTP thresholds can be dynamically matched, from now on we assumed that they are equal and static. Next, we wanted to investigate how the new nonlinear inhibitory plasticity rule adjusts the network response following a perturbation. Inspired by sensory deprivation experiments (Kirkwood et al., 1996; Philpot et al., 2003; Kuo and Dringenberg, 2009) or direct stimulation of input pathways (Huang et al., 1992; Abraham, 2008), we investigated the network response to perturbing the excitatory presynaptic input rate (Fig. 4A).

**Figure 4.**
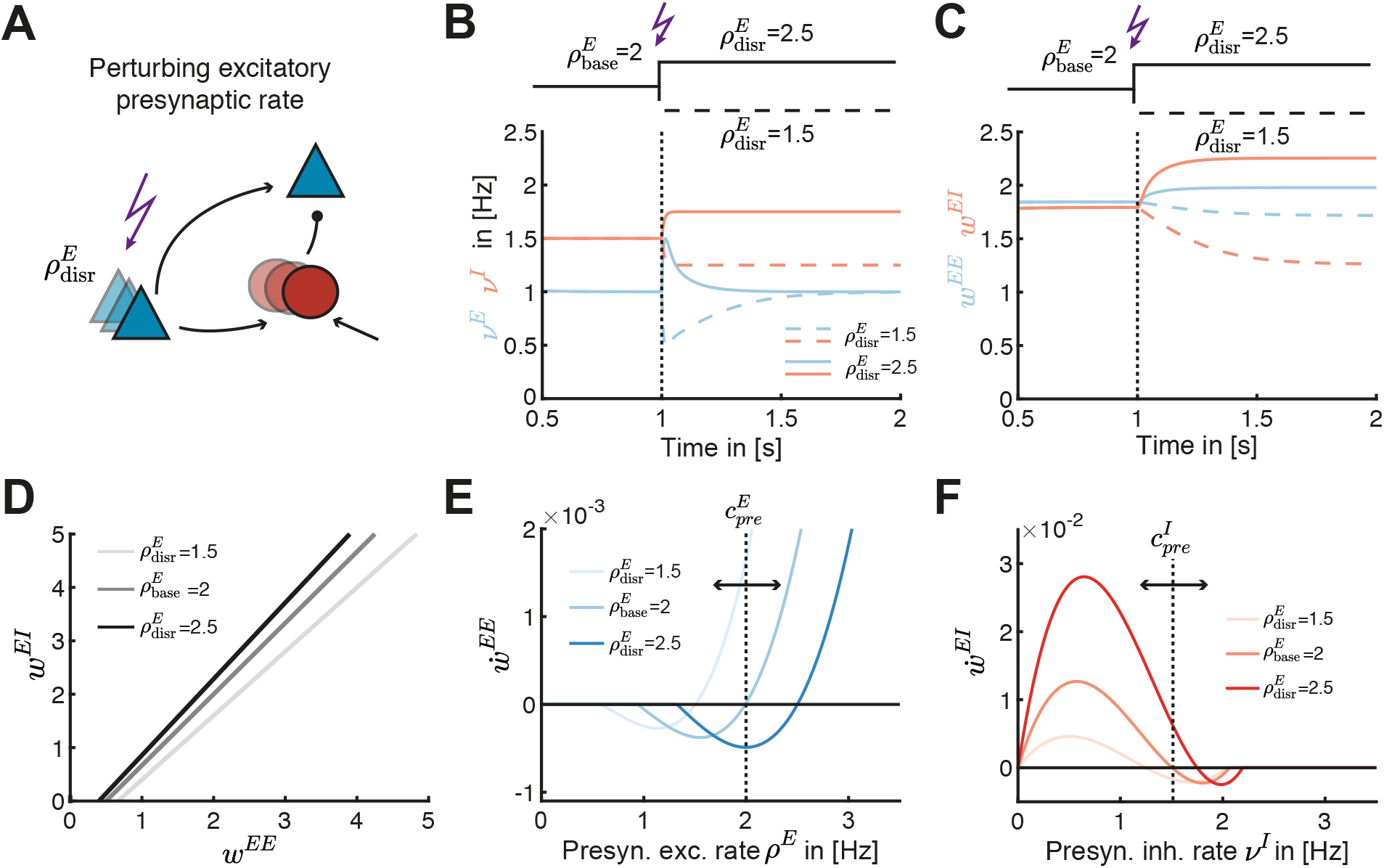
Nonlinear inhibitory plasticity can regulate the network response to perturbations. **A**. Schematic of perturbing the excitatory presynaptic rate in the inhibitory feedforward motif. **B**. Effect of increasing (solid lines, 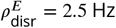) or decreasing (dashed lines, 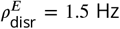) excitatory input rates from a baseline of 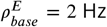 on excitatory (blue) and inhibitory (red) firing rates. **C**. Same as B but for the *w*^*EE*^ and *w*^*EI*^ weights. **D**. The line attractor for the baseline input 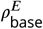 and two input perturbations 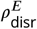. **E**. E-to-E weight change 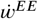 as a function of the presynaptic excitatory rate *ρ*^*E*^ for the baseline input 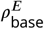 and for two input perturbations 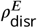. **F**. I-to-E weight change 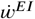 as a function of the inhibitory rate *v*^*I*^ for the baseline input 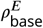 and for two input perturbations 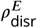.

Independent of the direction of the perturbation, we found that the novel inhibitory plasticity rule brings the excitatory postsynaptic rate back to the target rate (Fig. 4B). The inhibitory rate *v*^*I*^ also readjusts because the inhibitory population receives input from the perturbed excitatory population. But the new inhibitory rate is different than the rate before the perturbation (Fig. 4B). We found that a perturbation which decreases the excitatory input rate, leads to the depression of both type of weights *w*^*EE*^ and *w*^*EI*^ ; in contrast, a perturbation which increases the excitatory input rate leads to their potentiation (Fig. 4C). The response to these perturbations is consistent with previous experimental results. For example, it has been shown that input perturbations via sensory deprivation decrease inhibitory activity (Hengen et al., 2013; Kuhlman et al., 2013; Barnes et al., 2015). Specifically, sensory deprivation has been shown to depress inhibitory synaptic strengths, decrease in the number of inhibitory synapses (Chen et al., 2011; Keck et al., 2011; Chen et al., 2012; van Versendaal et al., 2012; Li et al., 2014) (but see (Maffei et al., 2006, 2010)) and depress excitatory synaptic strengths (Allen et al., 2003; Miska et al., 2018). At the same time, up-regulating activity has been shown to potentiate I-to-E synapses (Lourenço et al., 2014; Xue et al., 2014). Our framework can even predict the steady values of the E-to-E and I-to-E synaptic weights, as well as their ratio, by calculating the line attractor in the phase space of *w*^*EE*^ and *w*^*EI*^ weights as a function of the perturbed parameter (Fig. 4D).

Interestingly, we observed that this adjustment occurs by modulation of the presynaptic threshold between LTD and LTP for both excitatory and inhibitory plasticity. Decreasing the excitatory input rate decreases the excitatory presynaptic LTD/LTP threshold, hence limiting the range of presynaptic firing rates that generate depression. In contrast, we found that increasing the excitatory input rate increases the LTD/LTP threshold (Fig. 4E). Such a shift in the plasticity threshold for excitatory synapses has been measured in sensory deprivation experiments (Kirkwood et al., 1996; Philpot et al., 2003; Kuo and Dringenberg, 2009), and while restoring vision after sensory deprivation (Philpot et al., 2003; Cooper and Bear, 2012). Similarly to excitatory plasticity, perturbations in the excitatory input rate also shift the presynaptic threshold between LTD and LTP for inhibitory plasticity (Fig. 4F). Since there is no experimental evidence for this effect, we propose it as a prediction for the shift between LTD and LTP for I-to-E weights (*w*^*EI*^) in the presence of these perturbations.

In summary, our nonlinear inhibitory plasticity can adjust the network response and synaptic strengths to excitatory input rate perturbations, similar to experimental findings. We predict that this shift occurs by modulating the presynaptic LTD/LTP thresholds for both excitatory and inhibitory plasticity.

### The nonlinear inhibitory plasticity rule establishes a fixed excitatory to inhibitory weight ratio

We next wanted to investigate plausible functional roles of the newly proposed nonlinear inhibitory plasticity besides controlling excitatory and inhibitory firing rates and weights. Given our ability to calculate the steady states of the weights (Fig. 4D), we studied the ratio of E-to-E and I-to-E weights:

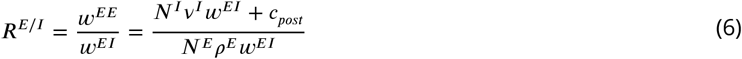

(Methods). For strong I-to-E weights *w*^*EI*^, the E/I ratio approximates to 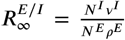 (Fig. 5A, inset; see Methods). Therefore, the E/I ratio is mainly determined by the ratio of excitatory and inhibitory input rates and the number of synapses. Paradoxically, the E/I ratio decreases as the presynaptic excitatory rate *ρ*^*E*^ increases (Fig. 5A; Eq. 6). This can be explained by considering that a higher excitatory input rate *ρ*^*E*^ generates more excitatory LTP (Fig. 1C), which needs to be counteracted by even more inhibitory LTP to stabilize the weights. Analytically, this corresponds to a line attractor with a steeper slope (Fig. 2D and Fig. 4D for increasing *ρ*^*E*^).

**Figure 5.**
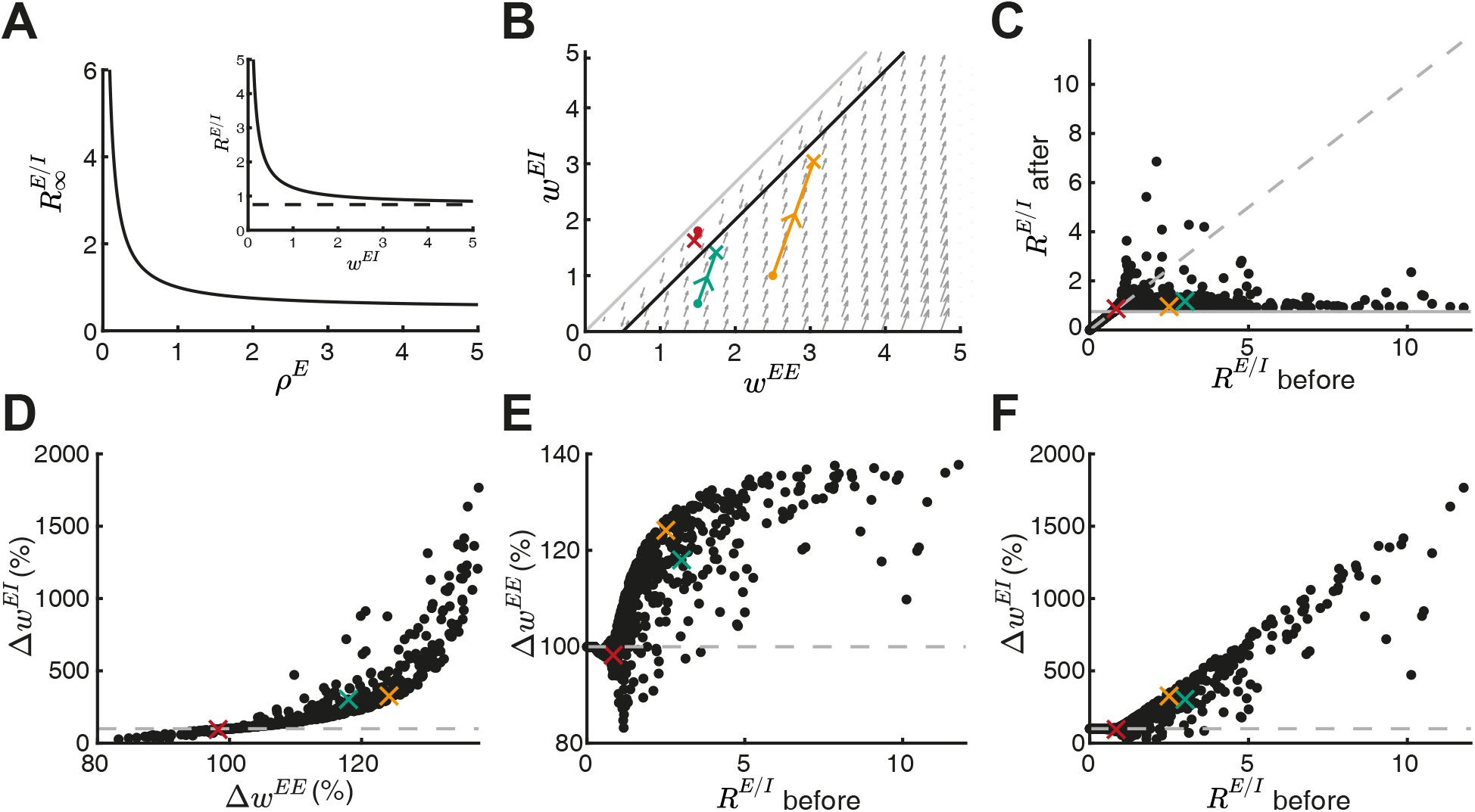
The nonlinear inhibitory plasticity rule keeps a fixed excitatory to inhibitory weight ratio. **A**. The steady state E/I weight ratio 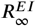 as a function of the presynaptic excitatory rate *ρ*^*E*^. Inset: ***R***^*E*/*I*^ approaches the steady state *N*^*I*^ *v*^*I*^ / *N*^*E*^ *ρ*^*E*^ (dashed line) for large I-to-E weights. **B-F** Based on a random initial weight configuration drawn from a uniform distribution in the range of [0, 3], excitatory and inhibitory plasticity was induced for 100 ms. Extreme initial E/I ratios (***R***^*E*/*I*^ before > 12) were excluded from the analysis. **B**. Phase portrait of the dynamics of E-to-E (*w*^*EE*^) and I-to-E (*w*^*EI*^) weights in the phase plane. Grey arrows indicate the direction of weight evolution over time, colored points represent three different weight initialization, 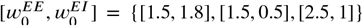, colored lines represents the weight evolution for each case and the cross marks the weights after plasticity induction. **C**. E/I ratio before and after plasticity induction. Crosses indicate examples in B. Grey dashed line indicates the identity line and grey line indicates 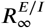 **D**. E-to-E weight change Δ*w*^*EE*^ versus I-to-E weight change Δ*w*^*EI*^ after plasticity induction in percent of initial synaptic weights. Dashed grey line indicates initial I-to-E weight strength and crosses indicate examples in B. **E**. E-to-E weight change Δ*w*^*EE*^ as a function of E/I ratio ***R***^*E*/*I*^ before plasticity in percent of initial weights. Dashed grey line indicates initial E-to-E weight strength and crosses indicate examples in B. **F**. Same as E but for I-to-E weight change Δ*w*^*EI*^.

Inspired by experiments (D’amour and Froemke, 2015), we evaluated the E/I ratio ***R***^*E*/*I*^ before and after inducing excitatory and inhibitory plasticity for multiple initial weight configurations (Fig. 5B,C; Methods). As predicted analytically (Fig. 5A), the E/I ratio after plasticity induction in these simulations approaches the set-point 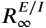 (Fig. 5C), matching experiments (D’amour and Froemke, 2015). E/I ratios far from the set-point before plasticity induction show the most drastic changes, with high postsynaptic firing rates resulting from dominant excitation needing to be overcome by fast and drastic weight changes by nonlinear inhibitory plasticity. Indeed, we observed that the I-to-E weights exhibit more change than E-to-E weights (Fig. 5D) in agreement with experiments (D’amour and Froemke, 2015). This suggests that nonlinear inhibitory plasticity plays a more prominent role than excitatory plasticity in establishing a fixed E/I ratio (Fig. 5E,F). With the linear inhibitory plasticity rule (Vogels et al., 2011), a fixed E/I ratio is only reached for initial weights which ultimately converge to the line attractor (Fig. 1E).

### Gating of receptive field formation via a disinhibitory signal

What functional implications does the proposed nonlinear inhibitory plasticity rule have on setting up network circuitry? Other than controlling excitatory and inhibitory rates and weights, here we wanted to examine if the nonlinear inhibitory plasticity rule can also enable flexible learning. Various forms of synaptic plasticity have been observed to support receptive field formation and generate selectivity to stimulus features in the developing cortex (Thompson et al., 2017). To investigate the function of interacting excitatory and inhibitory plasticity at the network level, we first extended the feedforward circuit motif to two independent pathways with pathway-specific inhibition (Fig. 6A). We found that perturbing the presynaptic excitatory rate of both inputs in opposite directions, decreasing for input 1 and increasing for input 2, differently shifts the input-specific excitatory presynaptic LTD/LTP thresholds and establishes different E/I ratios (Fig. 6B), both in agreement with experimental studies (Huang et al., 1992; Abraham and Bear, 1996). These results suggest that the control of E-to-E weight dynamics via nonlinear inhibitory plasticity is input-specific.

**Figure 6.**
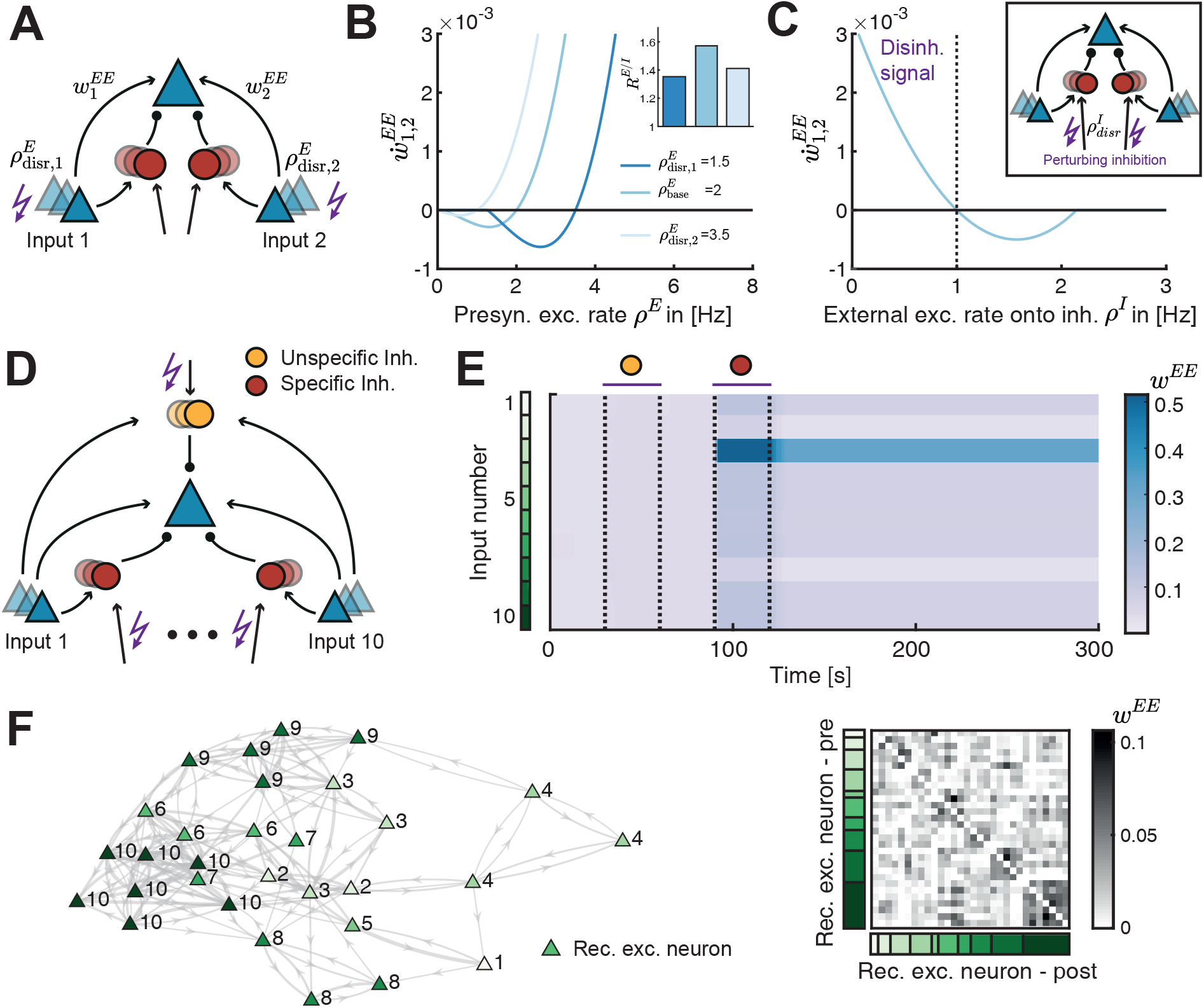
Gating of receptive field formation via a disinhibitory signal. **A**. Two independent inputs onto the same postsynaptic excitatory neuron. We perturb the presynaptic excitatory rate from input 1 or 2 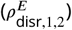. **B**. Plasticity curve of E-to-E weights for input 1 or 2 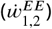 as a function of the presynaptic excitatory rate *ρ*^*E*^ for different input-specific perturbations 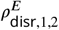. Inset: E/I weight ratio ***R***^*E*/*I*^ for different input-specific perturbations. **C**. Plasticity curve of E-to-E weights for input 1 and 2 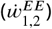 as a function of the external excitatory rate onto the inhibitory neurons *ρ*^*I*^, corresponding to a perturbation 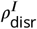 of the inhibitory populations. Perturbing *ρ*^*I*^ below 1 Hz (dashed line) is interpreted as a disinhibitory signal. Inset: We perturb the external excitatory rate onto the inhibitory neurons 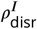. **D**. Ten independent inputs onto the same postsynaptic excitatory neuron with one inhibitory population unspecific to the input (yellow) and ten inhibitory populations each specific to one input (red). **E**. Evolution of excitatory weights over time. Purple bars indicate the time window for which disinhibition of either the unspecific (yellow) or all specific (red) inhibitory populations is applied (Methods). Input number color coded in green. **F**. Left: Network connectivity of recurrently connected excitatory neurons (triangles) after application of the disinhibitory signal. The number and the color indicates the input to which each neuron formed a receptive field (10 inputs in total). The thickness of the connection indicates the strength, only weights above 0.03 are shown. Right: Ordered recurrent E-to-E connectivity matrix. Input number color coded in green as in panel E.

Applying disinhibition by inhibiting the inhibitory population is a widely considered mechanism to ‘gate’ learning and plasticity (Froemke et al., 2007; Letzkus et al., 2011; Kuhlman et al., 2013). To test the potential of the circuit with nonlinear inhibitory plasticity to learn, we applied a disinhibitory signal by decreasing the external excitatory input onto the inhibitory populations. We found that this decreases the inhibitory input onto the postsynaptic neuron and potentiates E-to-E synapses, *w*^*EE*^ (Fig. 6C, *ρ*^*I*^ < 1). In contrast, increasing the input onto the inhibitory populations depresses E-to-E synapses (Fig. 6C, *ρ*^*I*^ > 1). Therefore, disinhibition via perturbation of the inhibitory neurons has the capacity to induce plasticity at E-to-E synapses and can gate excitatory plasticity.

How do the current results generalize to larger circuits with multiple independent inputs? In addition to pathway-specific inhibition, in this extended circuit we also introduced an unspecific inhibitory population (Fig. 6D). We presented different inputs to each pathway, corresponding to oriented bars in the visual cortex, or different single tone frequencies in the auditory cortex (Methods). We found that disinhibiting the unspecific inhibitory population does not selectively potentiate E-to-E weights, and hence does not generate competition among the different inputs. In contrast, disinhibiting all ten specific inhibitory populations strongly increases the E-to-E weights corresponding to only a subset of inputs, a process also called receptive field formation (Fig. 6E). This happens because the plasticity from the inhibitory population specific to the input stimulated at a given time counteracts any increases in the E-to-E weights from the same input. Hence, when the specific inhibitory population is inhibited, the increase of E-to-E weights is mostly balanced by unspecific I-to-E weights, leading to stimulus-specific differences in excitatory and inhibitory inputs and therefore to competition.

Finally, we implemented a network of 30 recurrently connected excitatory neurons where each neuron receives inputs from multiple inputs and an unspecific and a specific inhibitory population (Fig. 6F). In addition to the feedforward excitatory and inhibitory synapses, all recurrent E-to-E weights are also plastic. Similar as with a single postsynaptic neuron, we found that each of the excitatory neurons in the recurrent circuit forms a receptive field by becoming selective to one of the inputs (Fig. 6F, left; number next to the neuron). In addition, strong bidirectional connections form among recurrent excitatory neurons with similar receptive fields due to their correlated activity (Fig. 6F). This is consistent with strong bidirectional connectivity described in multiple experimental studies (Ko et al., 2011, 2013; Miller et al., 2014).

In summary, the newly proposed nonlinear inhibitory plasticity rule does not only ensure for stable synaptic weights and activity, but also enables the formation of feedforward and recurrent structures upon disinhibition which gates synaptic plasticity.

## Discussion

Hebbian excitatory synaptic plasticity is inherently unstable, requiring additional homeostatic mechanisms to control and stabilize excitatory-to-excitatory weight dynamics (Turrigiano and Nelson, 2004). Here, we proposed a novel form of inhibitory plasticity (Fig. 2), which can control excitatory and inhibitory firing rates and synaptic weights and enable stable and flexible learning of receptive fields in circuit models of the sensory cortex. We identified the dominance of inhibition over excitation (Eq. 4) and identical postsynaptic thresholds between LTD and LTP for excitatory and inhibitory plasticity as two key features for stabilization of weight dynamics in our model (compare Fig. 2A and Fig. 3A-C). However, the latter requirement can be relaxed with a suitable dynamic mechanism that enables self-adjusting of the plasticity thresholds in opposite directions for excitatory and inhibitory plasticity (Fig. 3D-F). This novel form of nonlinear inhibitory plasticity can also regulate the network response to perturbations of excitatory input rates (Fig. 4). A direct consequence of our inhibitory plasticity is the establishment of an E/I weight ratio set-point (Eq. 6), in agreement with experiments (D’amour and Froemke, 2015) (Fig. 5). Besides stability, the proposed form of inhibitory plasticity enables receptive field formation following disinhibition to input-specific inhibitory populations and in recurrent networks supports the formation of strong bidirectional connectivity among neurons with similar receptive fields (Fig. 6), suggesting a possible solution for the stability-flexibility problem.

### Inhibitory plasticity as a control mechanism of excitatory-to-excitatory weight dynamics

In the last decades, experimental studies have uncovered multiple possible mechanisms to counteract Hebbian runaway dynamics, including synaptic scaling (Turrigiano et al., 1998; Turrigiano, 2011), heterosynaptic plasticity (Lynch et al., 1977; Chistiakova et al., 2015), and intrinsic plasticity (Desai et al., 1999; Debanne et al., 2019). At the same time, computational studies have included multiple homeostatic mechanisms to stabilize rates and weight dynamics, including upper bounds on the E-to-E weights, normalization mechanisms (Oja, 1982; Miller and MacKay, 1994), and metaplastic changes of the plasticity function (Bienenstock et al., 1982; Yger and Gilson, 2015). However, the spatial and temporal scales for integrating Hebbian and homeostatic plasticity have remained an open question (Turrigiano, 2017; Zenke et al., 2017; Zenke and Gerstner, 2017).

In our study, we proposed a novel inhibitory plasticity rule at inhibitory-to-excitatory synapses which depends nonlinearly on the postsynaptic firing rate as a solution to the temporal paradox. While nonlinear excitatory plasticity rules have been identified in experimental studies (Kirkwood et al., 1996; Philpot et al., 2003; Cooper and Bear, 2012), less data is available for inhibitory plasticity. For example, presynaptic stimulation (hyperpolarization) and postsynaptic depolarization, have been shown to be required for inhibitory plasticity induction (Woodin et al., 2003; Chiu et al., 2018; Vickers et al., 2018; Udakis et al., 2020). Additionally, high-frequency stimulation of presynaptic input pathways has been shown to potentiate inhibitory synapses (Caillard et al., 1999; Shew et al., 2000; Mellor, 2018). Finally, the amount of inhibitory LTP has been shown to depend on the postsynaptic rate (Wang and Maffei, 2014). We designed our nonlinear inhibitory plasticity mechanism to be consistent with these findings: both, pre- and postsynaptic activity is necessary to induce inhibitory plasticity and the amount of LTP depends on the postsynaptic rate. Nonetheless, our rule is inconsistent with some experimental data which found no inhibitory plasticity for very high postsynaptic rates (Wang and Maffei, 2014).

### Inhibitory plasticity as a metaplastic mechanism

The ability of the proposed nonlinear inhibitory plasticity to control the sign and magnitude of excitatory plasticity resembles metaplasticity, i.e. a plasticity mechanism that is plastic itself (Yger and Gilson, 2015). We found that input perturbations modulate the excitatory presynaptic LTD/LTP threshold via a change of I-to-E weights and inhibitory rates consistent with metaplasticity (Fig. 4). Previous computational work has already suggested that a linear inhibitory plasticity rule can implement a metaplastic mechanism (Clopath et al., 2016). What mechanism underlies the sliding LTD/LTP threshold during the induction of plasticity is still an open question. Some experimental studies have suggested that inhibition can control sign and magnitude of excitatory plasticity (Paille et al., 2013; Vogels et al., 2013; Wang and Maffei, 2014; Hiratani and Fukai, 2017). Most intriguingly, it has been shown that application of gamma-Aminobutyric acid (GABA) can increase the excitatory LTD/LTP threshold, while blocking GABA can decrease the excitatory LTD/LTP threshold (Steele and Mauk, 1999), in close agreement with our findings (Fig. 1C).

The metaplasticity of excitatory plasticity was first suggested theoretically with the Bienenstock-Cooper-Munro (BCM) rule (Bienenstock et al., 1982), and was later confirmed in sensory deprivation and restoration experiments (Kirkwood et al., 1996; Philpot et al., 2003; Kuo and Dringenberg, 2009; Philpot et al., 2003; Cooper and Bear, 2012). In the BCM rule, the metaplastic mechanism is implemented by sliding LTD/LTP threshold dependent on the excitatory postsynaptic rate (Intrator and Cooper, 1992; Cooper et al., 2004). Higher (lower) postsynaptic rates lead to a higher (lower) postsynaptic LTD/LTP threshold making LTP (LTD) induction harder. An important difference in our model to the BCM rule is that the metaplastic sliding of the LTD/LTP threshold 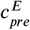 depends on the presynaptic excitatory rate (Fig. 1C), whereas the postsynaptic LTD/LTP threshold 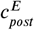 is fixed (except in Fig. 3D-F). This apparent difference can be resolved by assuming that homeostatic mechanisms operate at two different timescales: fast and slow. Slow homeostasis has been linked to a change in intrinsic excitability of neurons or synaptic scaling, for example, as during sliding of the postsynaptic LTD/LTP threshold as in the BCM rule (Keck et al., 2017). Fast homeostasis might be linked to disinhibition and inhibitory plasticity (Gainey and Feldman, 2017), and we suggest this is the case during sliding of the presynaptic LTD/LTP threshold mediated by inhibitory plasticity. Hence, it is plausible that both, presynaptic and postsynaptic metaplasticity exist in neuronal circuits. An advantage of homeostasis via inhibitory plasticity, rather than a direct influence on the E-to-E weights, might be that there is no interference with stored information in E-to-E connections.

### Key features of the nonlinear inhibitory plasticity rule

For the novel inhibitory plasticity rule to stabilize E-to-E weight dynamics, two key features need to be fulfilled. First, I-to-E weight changes need to be more ‘dominant’ than E-to-E weight changes (Fig. 2). More dominant means that I-to-E weights need to change with a higher magnitude at each time step compared to E-to-E weights, for all postsynaptic rates. If excitatory plasticity exceeds inhibitory plasticity for a certain postsynaptic rate as in the case of linear inhibitory plasticity, weight dynamics will be unstable (Fig. 1D-F). In our model, dominance of nonlinear inhibitory plasticity is guaranteed by the condition in Eq. 4, which involves relative number of synapses, presynaptic rates and plasticity timescales of excitation and inhibition to determine stability.

Second, matching the excitatory and inhibitory postsynaptic LTD/LTP thresholds, whereby excitatory and inhibitory synaptic change occur in the same direction for a given firing rate, is necessary for stable weight dynamics (Fig. 2A-C versus Fig. 3A-C). However, implementing a mechanism that dynamically shifts these thresholds in the opposite directions for excitatory vs. inhibitory plasticity based on experimental evidence (Keck et al., 2017), suggests that this match is not needed at all times. An interesting consequence from this dynamic threshold shift is a diversity of firing rates, which agrees with experimental data (Buzsáki and Mizuseki, 2014) and has recently been also achieved in different types of models (Pedrosa and Clopath, 2020; Agnes and Vogels, 2021).

We found that the new nonlinear inhibitory plasticity rule achieves an E/I ratio set-point (Fig. 5) in agreement with experimental data (D’amour and Froemke, 2015). We observed that inhibitory plasticity is the more dominant mechanism to achieve this. The dominance of inhibitory plasticity suggests a possible solution for the temporal paradox of integrating Hebbian excitatory plasticity and homeostasis (Zenke et al., 2017), eliminating the requirement for additional fast stabilizing mechanisms in our model. While the relative timescales of excitatory and inhibitory plasticity mechanisms remain an open question, most computational models agree on the need for faster inhibitory than excitatory plasticity dynamics (Sprekeler, 2017; Zenke et al., 2017).

### Functional implications of the nonlinear inhibitory plasticity rule

Our novel form of inhibitory plasticity leads to a fixed E/I balance, or more specifically to an E/I weight ratio setpoint (Fig. 5A,C and Eq. 6). This is consistent with several experimental studies which have suggested that inhibitory plasticity keeps an E/I ratio set-point (Froemke et al., 2007; Maffei and Turrigiano, 2008; Dorrn et al., 2010; House et al., 2011; Wang and Maffei, 2014; Xue et al., 2014; D’amour and Froemke, 2015; Adesnik, 2017; Field et al., 2020). For example, as our model would predict, some studies have found that the amount of inhibitory plasticity depends on how far the current E/I ratio is from the set point (Fig. 5F) (D’amour and Froemke, 2015; Aljadeff et al., 2019). Perturbing the excitatory input rate in our model as a model of sensory deprivation increases the E/I ratio (Fig. 5A), consistent with sensory deprivation experiments (Kuhlman et al., 2013; Li et al., 2014; Barnes et al., 2015; Miska et al., 2018). Despite the ability of the new nonlinear inhibitory plasticity rule to establish and maintain a fixed E/I balance, we acknowledge that there are various additional mechanisms that contribute, including heterosynaptic plasticity (Field et al., 2020).

The emergence of an E/I ratio set-point and the stabilization of rates driven by the novel inhibitory plasticity rule ensure a fixed E/I balance. E/I balance is usually more broadly defined as the proportionality of total excitatory and inhibitory input onto a neuron (Isaacson and Scanziani, 2011). In our model, once a fixed E/I balance is reached, there is no more synaptic plasticity and neurons fire at stable rates. To induce further weight changes, an additional gating signal is necessary that perturbs the E/I balance. In our model, there are three ways to gate plasticity: (1) directly changing the postsynaptic rate (Fig. 1B); (2) perturbing the excitatory input pathway (Fig. 4); and (3) perturbing the inhibitory population (Fig. 6C). The idea that inhibition gates excitatory plasticity is well-documented in the experimental literature (Dehorter et al., 2017; Hattori et al., 2017; Kripkee and Froemke, 2017).

Experimentally, both neuromodulation (Froemke et al., 2007; Froemke, 2015) and disinhibitory circuits (Letzkus et al., 2011, 2015; Wang and Yang, 2018; Williams and Holtmaat, 2019; Canto-Bustos et al., 2022) can directly control the activity of inhibitory neurons and lead to excitatory plasticity. Based on this, we investigated the gating of plasticity via a disinhibitory signal in the context of receptive field formation. While receptive field formation has already been demonstrated in multiple computational studies (Bienenstock et al., 1982; Luz and Shamir, 2012; Clopath et al., 2016), we propose that it can occur solely from the interaction of excitatory and inhibitory plasticity without any additional mechanism to induce competition among different inputs (Fig. 6D,E). Recurrently connecting multiple postsynaptic excitatory neurons and allowing the connections between them to be plastic leads to receptive field formation of each excitatory neuron in the recurrent circuit and the formation of strong bidirectional connectivity between neurons with similar receptive fields (Fig. 6F). This is in agreement with various experimental data (Ko et al., 2011, 2013; Miller et al., 2014; Lee et al., 2016) and has been previously achieved in models (Clopath et al., 2010; Litwin-Kumar and Doiron, 2014; Montangie et al., 2020).

Interestingly, we found that gating of receptive field formation via disinhibition depends on the specificity of the targeted inhibitory population to the inputs. While disinhibiting the unspecific population does not form receptive fields, disinhibiting all specific inhibitory populations induces competition between different inputs and forms receptive fields. If inhibitory plasticity counteracts excitatory plasticity in an input-specific way, no competition between input pathways can emerge because small biases in the E-to-E weights in one input are immediately balanced by I-to-E weights in the same input. Therefore, disrupting the specific inhibitory populations allows the strengthening of a subset of inputs. This result is similar to the findings of Clopath et al. (2016), where receptive field formation was shown to depend on the specificity of the inhibitory neurons.

The inhibitory populations in our model can be linked to the two main inhibitory neuron types in the cortex, somatostatin-expressing (SOM) and parvalbumin-expressing (PV) inhibitory interneurons. Specificity of the inhibitory neuron type to excitatory inputs can be interpreted as tuning of the inhibitory neurons to input features. In the visual (Ma et al., 2010; Cottam et al., 2013) and the auditory cortex (Li et al., 2015), tuning of SOM interneurons is sharper than PV interneurons, although conflicting evidence exists (Griffen and Maffei, 2014). Therefore, in our model the specific inhibitory neuron type could represent SOM interneurons while the unspecific inhibitory population could represent PV interneurons. Supporting this interpretation of SOM interneurons being the specific inhibitory population, experimental studies find that a suppression of SOM neurons gates excitatory plasticity (Chen et al., 2015; Hattori et al., 2017; Williams and Holtmaat, 2019),

### Predictions

We formulated our model with rate-based units not only because it enabled us to treat it analytically, but also because it led to an in-depth mechanistic understanding of the involved processes, allowing us to formulate experimentally testable predictions and making our model assumptions falsifiable. A main feature of our model is that inhibitory plasticity depends nonlinearly on the rate of the postsynaptic excitatory neuron. This can be tested experimentally by inducing inhibitory plasticity while varying the rate of an excitatory neuron and keeping the inhibitory input to this neuron constant. A second feature of our model is that excitatory and inhibitory plasticity have an identical postsynaptic LTD/LTP threshold. This could be tested by inducing plasticity at excitatory and inhibitory pathways onto the same excitatory neuron, and measuring the LTD/LTP thresholds as a function of the rate of that neuron.

Based on the perturbation experiment (Fig. 4), we can formulate multiple predictions. First, we hypothesize that the mechanism behind the metaplastic mechanism is a change in the level of inhibition (see Fig. 1C, Fig. 4E). Therefore, blocking inhibitory plasticity experimentally should also disrupt the metaplastic mechanism. Second, we predict that the shape of inhibitory plasticity as a function of the inhibitory rate is reversed compared to excitatory plasticity, and perturbations of the excitatory input lead to specific metaplastic changes of inhibitory plasticity. Decreasing the excitatory input should lower the inhibitory LTD/LTP threshold as a function of the presynaptic inhibitory rate and decrease the inhibitory LTP magnitude (Fig. 4F). Third, following from the dependence of the line of stable fixed point on several model parameters (Fig. 2C and Eq. 5), especially on the excitatory input rate (Fig. 4D), we hypothesize that different E/I ratios can be achieved following input perturbations. Decreasing the excitatory input rate should lead to higher E/I ratios, while increasing it to lower E/I ratios.

The new rule suggests a new functional role of inhibitory plasticity, namely controlling E-to-E weight dynamics. Therefore, we extend previously studied roles of inhibitory plasticity, which include the stabilization of excitatory rates (Vogels et al., 2011; Sprekeler, 2017), decorrelation of neuronal responses (Duarte and Morrison, 2014), preventing winner-take-all mechanisms in networks with multiple stable states (Litwin-Kumar and Doiron, 2014) or generating differences among novel versus familiar stimuli (Schulz et al., 2021). Recent computational studies also include novel ways of inhibitory influence, like presynaptic inhibition via GABA spillover (Naumann and Sprekeler, 2020) or an input-dependent inhibitory plasticity mechanism (Kaleb et al., 2021).

Our model included a single type of inhibitory plasticity. Yet, recent studies have found that cortical circuits have abundance of different inhibitory interneuron types and that inhibitory plasticity depends on the inhibitory neuron type (Chiu et al., 2018; Vickers et al., 2018; Udakis et al., 2020; Lagzi et al., 2021). Our result on inhibitory population-dependent effects in gating receptive field formation suggests that subtype-specific plasticity rules might have non-trivial influences on the network, as some recent models have proposed (Agnes et al., 2020; Lagzi et al., 2021). Furthermore, other homeostatic mechanisms will influence the stability of weight dynamics, E/I ratio set-points and the effect different perturbations have on the network dynamics.

### Conclusion

Taken together, our study proposed a novel form of nonlinear inhibitory plasticity which can achieve stable firing rates and synaptic weights without the need for any additional homeostatic mechanisms. Moreover, our proposed plasticity is fast, and hence could provide a solution to the temporal paradox problem because it can counteract fast Hebbian excitatory plasticity. Functionally, our proposed inhibitory plasticity can establish and maintain a fixed E/I ratio set-point. At this set-point, no synaptic plasticity is induced, i.e. plasticity is “off”. Perturbing the postsynaptic firing rate, e.g. via disinhibition, can act as a gate, turning plasticity “on”. This enables the competition among input streams leading to receptive field formation in feedforward and recurrent circuits. Therefore, our nonlinear inhibitory plasticity mechanism provides a solution to the stability-flexibility challenge.

## Methods

### Rate-based model

We studied rate-based neurons to allow us to analytically investigate the dynamics of firing rates and synaptic weights in the model. In the feedforward motif (Fig. 1A), we considered a network consisting of one excitatory postsynaptic population with a linear threshold transfer function and firing rate *v*^*E*^, receiving input *N*^*E*^ presynaptic excitatory populations with firing rates *ρ*^*E*^ through synapses with weights *w*^*EE*^, and *N*^*I*^ presynaptic inhibitory populations with firing rates *v*^*I*^ through synapses with weights *w*^*EI*^ :

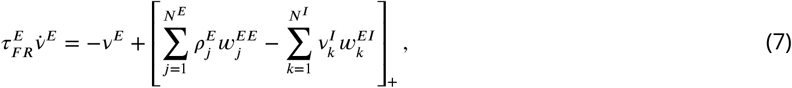

where []_+_ denotes a rectification that sets negative values to zero. The inhibitory neurons also follow a similar dynamics and are driven by the same *N*^*E*^ presynaptic excitatory populations with firing rates *ρ*^*E*^ through synapses with weights *w*^*IE*^ and additional external input with firing rate *ρ*^*I*^,

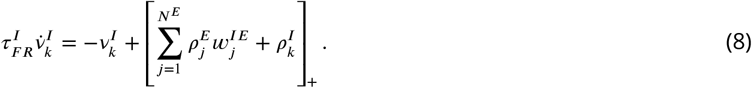

Here, 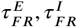 denote the time constants of the firing rate dynamics. All parameters are given in Table 1. The synaptic weights, *w*^*EE*^ and *w*^*EI*^ are plastic according to different plasticity rules (see below). For simplicity, we do not use subscripts for neuron identity and interpret all variables as mean values and hence can denote the total excitatory input to the postsynaptic neuron as *N*^*E*^ *ρ*^*E*^ *w*^*EE*^ and the total inhibitory input as *N*^*I*^ *v*^*I*^ *w*^*EI*^. In the mean-field sense, also the number of neurons can be traded-off with the rates or the synaptic weights, hence we assume *N*^*E*^ = *N*^*I*^ = 1 (Table 1).

### Rate-based plasticity

For the plasticity of E-to-E synaptic weights *w*^*EE*^, we used a learning rule that depends nonlinearly on the postsynaptic rate *v*^*E*^ (Fig. 1B) (Kirkwood et al., 1996; Philpot et al., 2003; Cooper and Bear, 2012):

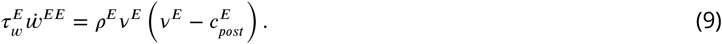

Here, 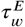 is the timescale of excitatory plasticity which is much longer than the timescale of the neuronal dynamics. The plasticity changes sign at the ‘postsynaptic LTD/LTP threshold’, 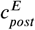.

For the plasticity of I-to-E synaptic weights *w*^*EI*^, we used two learning rules. First, we used an inhibitory plasticity rule common in computational models (Vogels et al., 2011; Clopath et al., 2016), which depends linearly on the postsynaptic rate *v*^*E*^ (Fig. 1D, 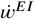):

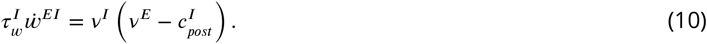

Here, 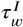 denotes the timescale of inhibitory plasticity which again is much longer than the timescale of the neuronal dynamics. As for excitatory plasticity, inhibitory plasticity changes from LTD to LTP at the ‘inhibitory postsynaptic LTD/LTP threshold’, 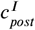, which sets the ‘target rate’ of the postsynaptic neuron (Vogels et al., 2011). In our paper, we proposed a novel inhibitory plasticity rule, which also depends nonlinearly on postsynaptic excitatory activity just like excitatory plasticity (Fig. 2A):

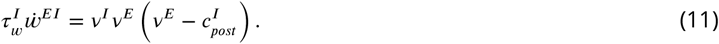

For both inhibitory plasticity rules, we assumed that the excitatory and inhibitory thresholds are matched 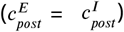 to prevent excitatory and inhibitory plasticity pushing the postsynaptic excitatory neuron towards two different firing rates. The except for this was the dynamic mechanism for thresholds adjustment in Fig. 3.

#### LTD/LTP plasticity thresholds

As can be see in the equations for excitatory and inhibitory plasticity, the postsynaptic LTD/LTP thresholds, which determine the sign of plasticity as a function of postsynaptic excitatory activity, are fixed. However, in the main text we also introduce the concept of a presynaptic LTP/LTD thresholds, defined as the presynaptic excitatory rate at which no synaptic plasticity is induced. We consider *v*^*E*^ at steady state (*v*^*E*^ = [*N*^*E*^ *ρ*^*E*^ *w*^*EE*^ −*N*^*I*^ *v*^*I*^ *w*^*EI*^]_+_) and assume that the dynamics of the rates is in the region where the transfer function is linear. Therefore, we can drop the linear rectifier and solve for *ρ*^*E*^ at which Eq. 1 is zero. We derive the presynaptic LTD/LTP threshold as:

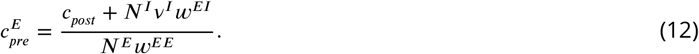

#### Stability analysis

To investigate the stability of the weights, we first calculated the nullclines, where we assumed that the postsynaptic excitatory rate is at steady state *v*^*E*^ = [*N*^*E*^ *ρ*^*E*^ *w*^*EE*^ − *N*^*I*^ *v*^*I*^ *w*^*EI*^]_+_. By setting Eqs. 9 and 11 to zero and dropping the linear rectifier, i.e. *v*^*E*^ = *N*^*E*^ *ρ*^*E*^ *w*^*EE*^ − *N*^*I*^ *v*^*I*^ *w*^*EI*^, we can write

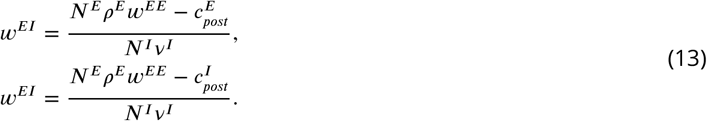

We see that the two equations are identical if 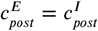. Therefore, only for identical LTD/LTP thresholds 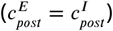 a line of fixed points emerges. The fixed points are 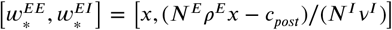, where we require that *x* ≥ *c*_*post*_/ (*N*^*E*^ *ρ*^*E*^) to avoid negative weights. To calculate the stability of the line of fixed points, we calculate the eigenvalues. We can rewrite Eqs. 9 and 11, as

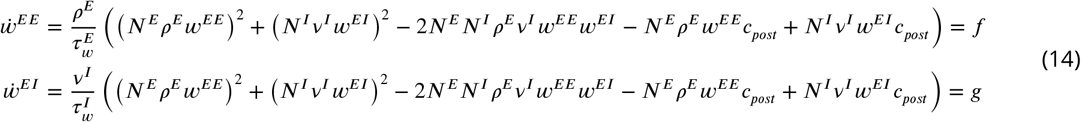

where we drop the linear rectifier by assuming that the dynamics of the rates is in the region where the transfer function is linear. We find that the entries of the Jacobian matrix at the fixed points are

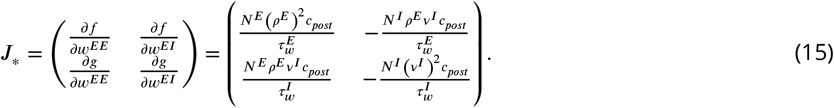

The trace of the Jacobian is 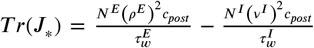 and the determinant is zero *Det* (*J*_*∗*_) = 0, therefore we find that the eigenvalues are:

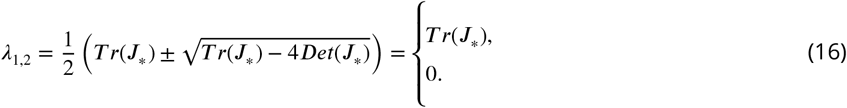

This means that if *T r*(*J*_*∗*_) < 0, the system is stable. Reordering this condition leads to the stability condition derived in the main text as Eq. 4. By reordering the terms in the nullclines given in Eq. 13, we derive the line attractor equation as given in the main text in Eq. 5.

The nonlinear excitatory and inhibitory plasticity rules have a fixed point when the postsynaptic excitatory firing rate is *v*^*E*^ = 0 Hz. Therefore, in the phase plane of *w*^*EE*^ and *w*^*EI*^ weights there is a region where the total inhibitory input is larger than the total excitatory input, *N*^*E*^ *ρ*^*E*^ *w*^*EE*^ < *N*^*I*^ *v*^*I*^ *w*^*EI*^, resulting in no postsynaptic firing (Fig. 2B, above grey line). The line equation separating the space with and without weight dynamics is

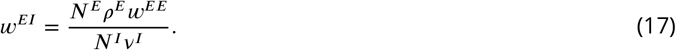

In the case of the linear inhibitory plasticity rule, stability depends on the initial weights. The line which separates stable from unstable initial weights can be calculated by taking the ratio of Eq. 9 and Eq. 10 and equating that to the slope of the line attractor (Eq. 5):

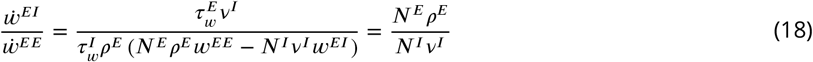

which leads to

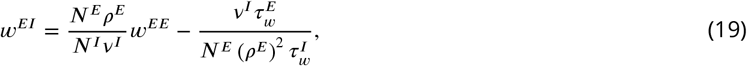

which is the equation of the dashed line in Fig. 1E.

### Dynamic threshold matching

The equations for the dynamics postsynaptic LTD/LTP thresholds in Fig. 3D-F are

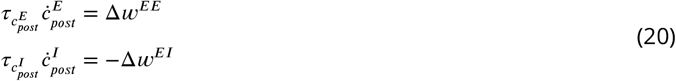

and therefore 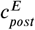 increases (decreases) if the postsynaptic neuron fires at 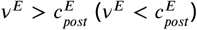 and 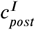 decreases (increases) if the postsynaptic neuron fires at 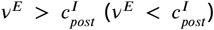. The amount of increase or decrease of the postsynaptic thresholds is scaled by the amount of plasticity induction, and we used a timescale of 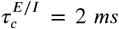, which is faster than the timescale of excitatory and inhibitory plasticity (Table 1).

### E/I ratio

We can calculate the E/I weight ratio ***R***^*E*/*I*^ in Eq. 6 by rewriting Eq. 13 and dividing one of the nullclines by *w*^*EI*^. For large weights, or in mathematical terms for *w*^*EI*^ → ∞, the E/I ratio becomes 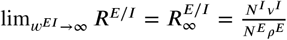.

To calculate the E/I ratio ***R***^*E*/*I*^, we take the solution for one nullcline from Eq. 13 (since 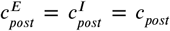 both are equivalent) and re-order the terms to reach Eq. 6. In the feedforward circuit (Fig. 1A), it follows

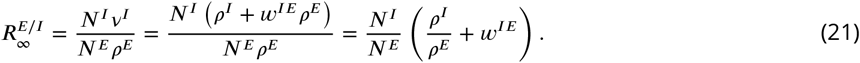

For the assumption *N*^*E*^ = *N*^*I*^ it follows that for larger excitatory input rate *ρ*^*E*^ the E/I ratio reaches a set-point at 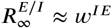 (see Fig. 5A, where *w*^*IE*^ = 0.5). Therefore, the E/I ratio has a lower bound which depends on the strength of the connection from the excitatory-to-inhibitory population.

In Fig. 5, we link our model to the experimental findings in D’amour and Froemke (2015). In D’amour and Froemke (2015), the authors induce plasticity with a spike-pairing protocol, in which pre-post spikes elicit excitatory LTP while post-pre spikes elicit LTD. Inhibitory LTP was induced for short time differences between the pre- and postsynaptic spikes (independent of the order of the spikes) and inhibitory LTD for longer time differences of the spike pairs. Since in the experiments the presynaptic stimulation was done with a stimulation electrode, the excitatory and inhibitory inputs did not have to be functionally related. In the model, we randomly drew initial E-to-E and I-to-E weights and induced plasticity for a limited amount of time (100ms) based on the rate-based plasticity rules (Eqs. 9 and 11).

### Gating of receptive field formation and recurrent clustering

Here, we explore a feedforward network with multiple inputs and two inhibitory neuron populations (Fig. 6C). To form receptive fields, we provide a random patterned input to the network. We define a pattern to mean an increase in the input rate to 4 Hz for 100 ms of four neurons. We then disinhibit the postsynaptic neurons by inhibiting either the total unspecific or specific inhibitory populations for 60 s by inducing an inhibitory input of 2 Hz onto the respective inhibitory neuron population. We model the release of disinhibition for the specific inhibitory population as slow and gradual over a time course of 100 s to avoid complete silencing of the postsynaptic excitatory neurons. We also note that here we used instantaneous integrators, i.e. 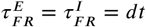 (Table 2), because we only wanted to focus on the interaction of excitatory and inhibitory plasticity in the model, though see (Gjorgjieva et al., 2016).

**Table 2.**
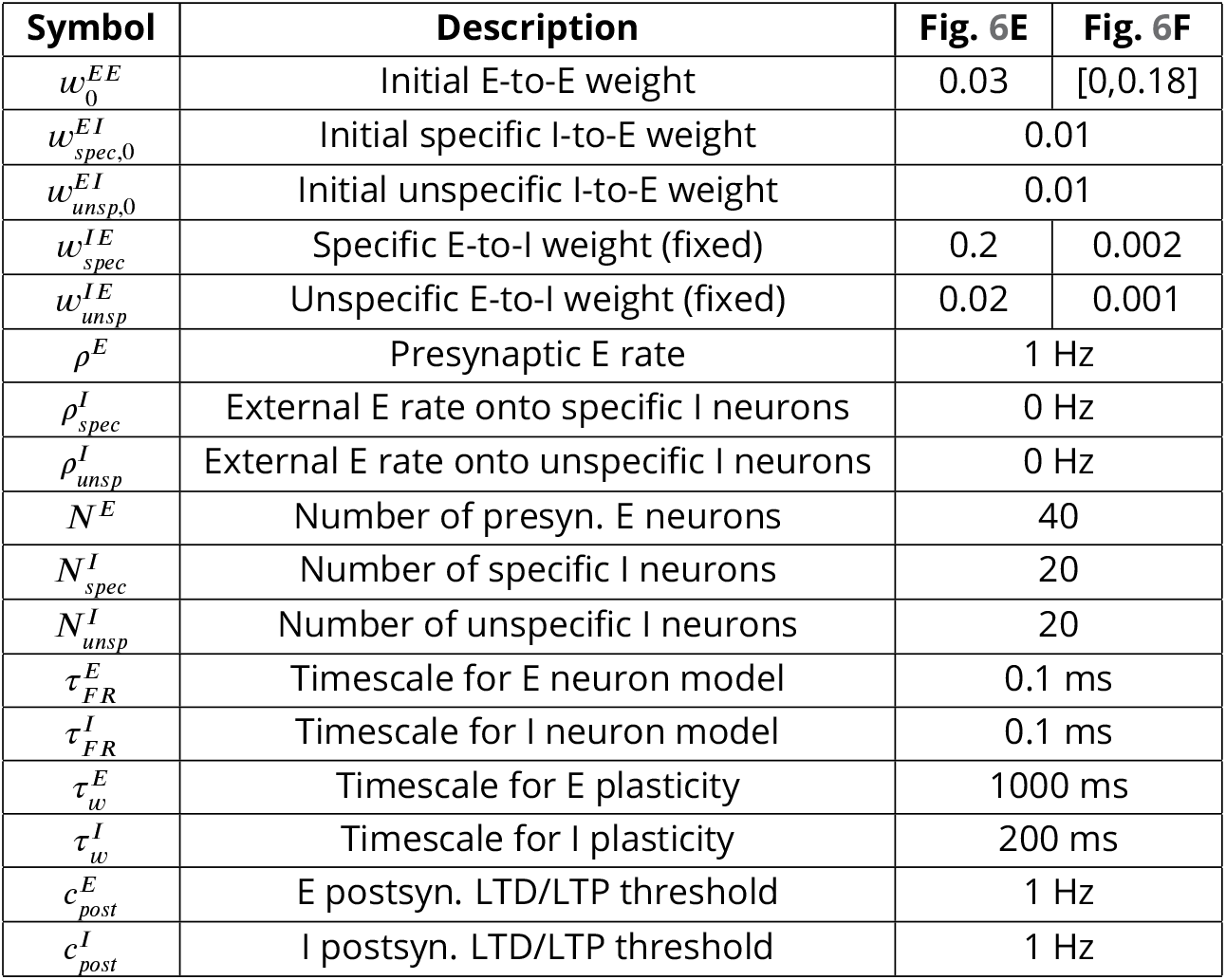
Parameter values for Fig. 6E,F.

For the recurrent circuit, we connected recurrently 30 postsynaptic neurons with feedforward circuits with specific and unspecific inhibition as described above (see also Fig. 6D,E) using an initial weight of 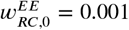. In addition to feedforward excitatory and inhibitory weights, also recurrent excitatory weights were plastic based on the plasticity mechanism of Eq. 9. We allowed the input patterns to each of the recurrent excitatory neuron to be correlated. Initial recurrent excitatory weights were randomly drawn from the interval [0,0.18]. To get the input to which recurrent neurons formed a receptive field to, we calculated the mean weight per input pattern and chose the maximum of those to be the input the neurons formed a receptive field to. The clustering graph in Fig. 6F (left) was done with the digraph function in Matlab.

The simulations were performed using Matlab programming language. Euler integration was implemented using a time step of 0.1 ms. Code implementing our model is available here: https://github.com/comp-neural-circuits/Nonlinear-inhibitory-plasticity.

## Acknowledgements

CM and JG thank the Max Planck Society for funding and a NARSAD Young Investigator Grant from the Brain and Behavior Research Foundation to JG. We also thank the Deutsche Forschungsgemeinschaft (DFG) for funding through the Collaborative Research Centre (CRC) 1080. We thank all members of the ‘Computation in Neural Circuits’ group for useful discussions and comments on the manuscript.

## Supplementary Material

**Figure S1.**
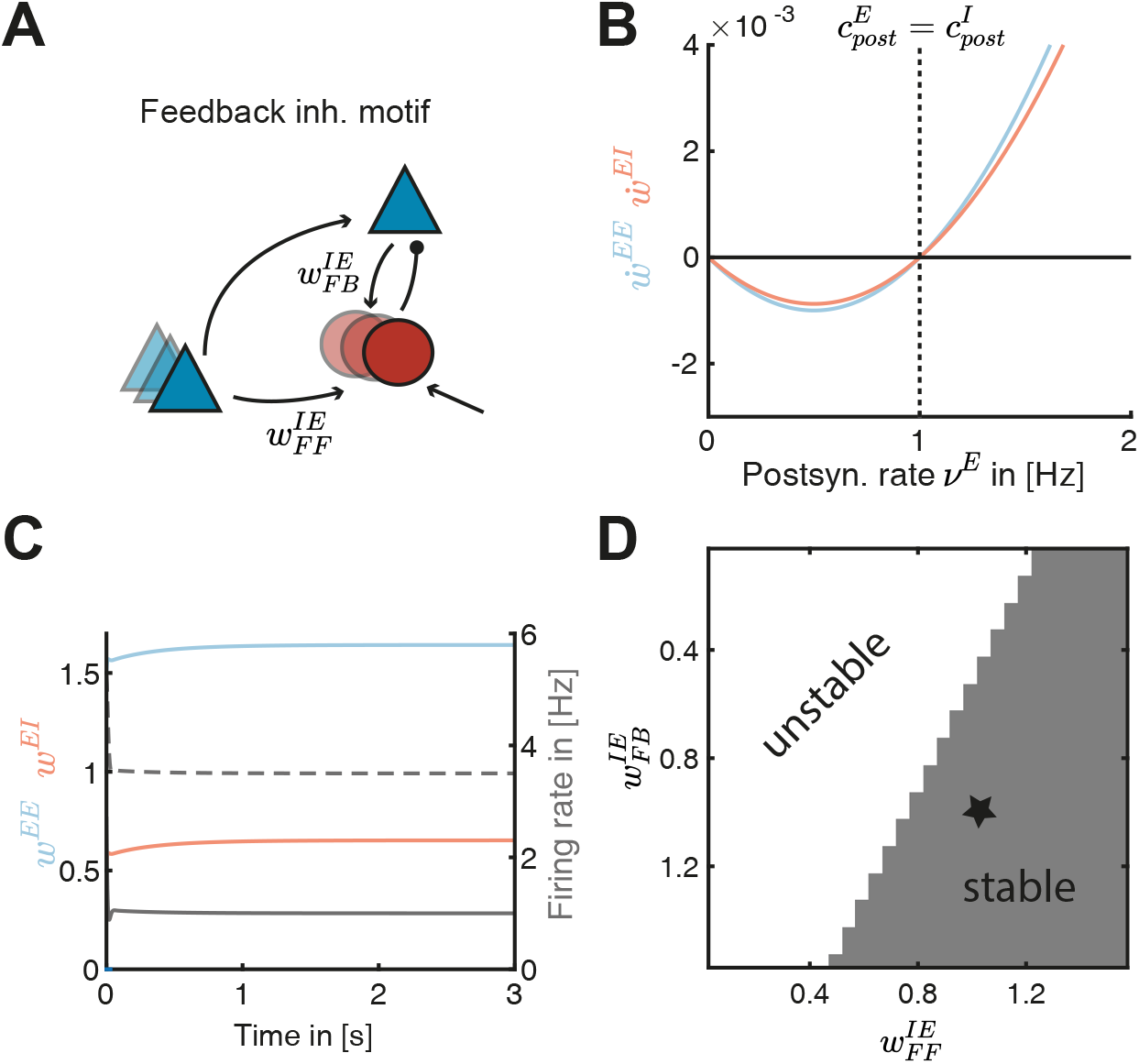
Feedback inhibitory motif leads to additional stability. **A**. Schematic of the feedback inhibitory motif. The inhibitory population receives input from the presynaptic excitatory population with weight strength 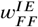 and the excitatory post-synaptic neuron with weight strength 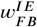. **B**. Plasticity of E-to-E (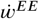, blue) and I-to-E (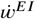, red) weights as a function of the postsynaptic rate *v*^*E*^. The excitatory and inhibitory LTD/LTP thresholds are identical 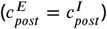. **C**. E-to-E (*w*^*EE*^, blue) and I-to-E (*w*^*EI*^, red) and rate dynamics of the postsynaptic (grey line) and the inhibitory population (grey dashed line) as a function of time. **D**. Stability of weight dynamics as a function of the excitatory-to-inhibitory weights 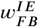 and 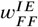. Star indicates the values shown in panel C.

